# The immunomodulatory properties of the HDAC6 inhibitor ACY241 supports robust anti-tumor response in NSCLC when coupled with the chemotherapy drug Oxaliplatin

**DOI:** 10.1101/2021.10.01.462824

**Authors:** Arup Bag, Andrew Schultz, Saloni Bhimani, William Dominguez, Ling Cen, Dennis Adeegbe

## Abstract

**Background:** Durable treatments that benefit a wide pool of patients remain elusive for Non-small cell lung cancer (NSCLC). The success of immunotherapy in a subset of NSCLC patients highlights the potential contribution of immune response to anti-tumor immunity while underscoring a need for broadly applicable therapeutic strategies. HDAC inhibitors are a promising class of drugs whose immunomodulatory properties are now being appreciated. In the present study, we evaluated the effects of the HDAC6 inhibitor, ACY241 on lung tumor immune compartment with the goal of understanding the scope of its immunomodulatory properties and its therapeutic potential in combination with Oxaliplatin.

**Methods:** Lung adenocarcinoma-bearing mice were treated with ACY241 or vehicle after which the proportions and phenotype of tumor-associated T cells and macrophages were evaluated by comprehensive flow cytometric analysis. Bulk RNA-sequencing was also conducted on both cellular subsets to interrogate the transcriptomic changes associated with ACY241 treatment relative to vehicle controls. *In vivo* drug efficacy study was performed by administration of ACY241 and/or Oxaliplatin and assessing tumor growth and survival of tumor-bearing mice. *Ex vivo* functional studies was performed to assess tumor-associated T cell effector function as it correlates with measured outcomes.

**Results:** We demonstrate that ACY241 promotes increased presence of T and NK cells in the lung tumors of treated mice. The tumor-associated T cells under ACY241 treatment displayed enhanced activation, proliferation, and effector profile. In addition, tumor-associated macrophages exhibited increased expression of MHC and co-stimulatory molecules while expression of inhibitory ligands were reduced. RNA-sequencing of both tumor-associated T cells and macrophages revealed significant genomic changes in both subsets that is consistent with ACY241-mediated enhancement of immune priming. These broad immunomodulatory properties of ACY241 were associated with significantly enhanced tumor-associated T cell effector functionality, robust anti-tumor response, and significantly prolonged survival of NSCLC-bearing mice when combined with the chemotherapy drug Oxaliplatin.

**Conclusion:** Collectively, our studies highlight the broad immunomodulatory effect of ACY241 as a promising HDAC6 inhibitor which coupled with Oxaliplatin promotes robust therapeutic outcomes in a pre-clinical model of NSCLC, providing compelling rationale for the clinical testing of this novel combinatorial regimen in NSCLC.

## Introduction

Non-small cell lung cancer (NSCLC) which accounts for over 80% of all lung cancers is a leading cause of cancer-related deaths worldwide. The success of immunotherapy drugs evidenced by the durable response achieved in a subset of NSCLC patients highlight the inherent potential of immune cells to orchestrate effective anti-tumor response. PD-1 blockade for instance promotes objective response in about 20% of patients ^1^ with higher tumor-expressed PD-L1 ^2^ or mutational burden ^3 4^ correlating with higher overall response rate. Targeting CTLA-4 also supports improved anti-tumor response as anti-CTLA-4 plus anti-PD-1 treatment was associated with an objective response rate of 35.9% compared to 30% with chemotherapy ^5^. While promising, these outcomes still serve as a reminder for the unmet clinical need for drugs that could benefit a wider pool of patients.

Histone deacetylase (HDAC) inhibitors are increasingly being evaluated in oncologic applications as promising therapeutic agents for cancer therapy in part due to their cytotoxic and cytostatic effects on tumor cells ^6 7^. The pan-HDAC inhibitors Panobinostat and Vorinostat are FDA-approved agents for multiple myeloma and cutaneous T-cell lymphoma, respectively ^8-10^. In recent years, research into the utility of these and other pan-HDAC inhibitors in solid tumors has increased considerably ^6 7 11-13^. However, due to potential toxicity concerns, more selective HDAC inhibitors are being increasingly considered as alternatives to pan-HDAC inhibitors based on expanding knowledge of the roles of various HDAC proteins to cancer biology. Among the selective HDAC inhibitors currently being explored are those targeting members of the class I (1, 2, 3, 8) and II (4, 5, 6, 7, 9, 10) HDACs which regulate various aspects of cancer biology including, proliferation, survival, metabolism, metastasis, autophagy, and apoptosis ^13-15^.

Despite their direct tumor cell toxicity, the effects of pan and isozyme selective HDAC inhibitors on immune cells is an important issue of consideration given the crucial contribution of immune cells to shaping the course of tumor progression. A number of HDAC inhibitors have been described to possess properties that have direct relevance to immune response in cancer. For example, panobinostat has been shown to promote upregulation of PD-1 ligands in melanoma, an effect that facilitated augmented anti-tumor response under anti-PD-1 therapy ^16^. Trichostatin A and panobinostat enhanced the expression of MHC class I and tumor-related antigens in tumor cells in a murine melanoma model thereby enhancing their immunogenicity ^17-19^. With respect to immune cells, HDAC inhibitors can exert modulatory properties ^20-22^. In alignment with this notion, romidepsin enhanced expression of T cell recruiting chemokines by tumor cells facilitating T cell infiltration into lung tumors in a mouse model of lung cancer ^22^. Understanding the nature and scope of immunomodulatory capabilities of HDAC inhibitors is no doubt paramount to their utility as suitable partner agents in novel combinatorial drug regimens. In this regard, the present study evaluated the effects of ACY241, an orally available HDAC6 inhibitor which while structurally similar to ricolinostat, is more potent with respect to its selectivity for HDAC6 (IC50 of 2.6 nM versus 5 nM in addition to having improved solubility properties ^23 24^.

Although its immune modulating properties has been described in a number of solid and hematologic cancers ^23-25^, its immunoregulatory effect and therapeutic potential in clinically relevant models of non-small cancer has not been substantiated. Given its favorable drug profile ^24^, the present study was undertaken to determine the spectrum of ACY241’s effect on tumor-infiltrating T cells and macrophages in a pre-clinical mouse model of NSCLC with emphasis on molecular underpinnings of drug effect and its potential utility as a partner agent in a rationally-selected combinatorial drug regimen for NSCLC therapy.

## Material and Methods

### Mice

All breeding and treatment experiments were performed with the approval of Moffitt Cancer Center Animal Care and Use Committee. Mice were maintained under specific-pathogen-free conditions in an AAALAC-accredited facility. Genetically engineered mice Kras^+/LSL-G12D^; Trp53^L/L^ (KP) have been previously described ^26^ For lung tumor induction, mice received 1 × 10^6^ CFU Cre-encoding adenovirus intranasally at 5–6 weeks of age. Tumor formation was evaluated by MRI with BioSpec USR70/30 horizontal bore system (Bruker). Tumor burden was quantified from the MRI images with 3D Slicer software. All mice were maintained under pathogen-free conditions in accordance with institutional guidelines for animal welfare

### *In vivo* Treatment Studies

After MRI-confirmation of tumors, mice with established tumors (150-200mm^3^) were randomly assigned to treatment groups. For short-term treatment studies involving genomic profiling by RNA-sequencing, KP mice were treated with ACY241 (25 mg/kg body weight) 3x/week for two weeks. Control mice received vehicle (5% DMSO in 10% 2-hydroxylpropyl β-cyclodextrin). For long-term drug efficacy studies, ACY241 was administered 3x/week and Oxaliplatin (10 mg/kg body weight) was administered 1x/week in either the single agent or the combination groups for 6 weeks after which they were monitored until clinical endpoint. For mice with OVA-expressing tumors, KP mice were inoculated with B6 OVA 10103 F LT1 OVA-PGK tumor cell line (kind gift from Dr. Tyler Jacks, MIT). After tumor establishment as documented by MRI, mice were assigned into vehicle or ACY241 treatment groups. ACY241 or vehicle was administered 3x/week for 2 weeks after which tumors were harvested for flow cytometry and ImageStream analysis.

### Immune profiling with multicolor flow cytometry

Tumor nodules collected from the lungs of KP mice at time of analysis were cut into 1mm pieces and placed in mouse tumor dissociation kit (Miltenyi Biotec) for 45 minutes at 37°C. After digestion, cells were passed through a 70 µm strainer to remove cellular debris and treated with ACK Lysing Buffer (Life Technologies). Cells were resuspended in 1X PBS, counted, and 1×10^6^ cells per sample were stained for flow cytometry. Briefly, cells were first incubated with LIVE/DEAD Fixable Aqua Dead Cell Staining Kit (Invitrogen, Carlsbad, CA) for 20 minutes at 4 °C followed by two washes with FACS buffer (1X PBS+ 2% FBS). FcR block was performed by incubating cells with purified anti-mouse CD16/32 (BioLegend) for 5 minutes at room temperature followed by surface staining with antibody mixture at 4^0^ in the dark for 20 minutes. Cells were then washed twice and resuspended in FACS buffer. Intracellular staining was performed for Foxp3 and other nuclear proteins using the Foxp3 staining kit (eBioscience, Santa Clara, CA) according to manufacturer’s instructions. After two washes, samples were resuspended in FACS buffer before acquisition using the BD FACSymphony (BD Biosciences) and analyzed with FlowJo software (Treestar). Gating strategy for FACS analyses is shown in figure S9 (supplemental file 2).

### Cell purifications and *Ex vivo* T cell activation assay

T cells from lung tumors of treated mice were isolated from single cell suspensions using CD90.2 Magnetic beads (Miltenyi) according to the manufacturer’s protocol. 1 × 10^6^ isolated T cells were then stimulated at 37°C with eBioscience™ Cell Stimulation Cocktail plus protein transport inhibitors (Invitrogen) for 6 hours. Cells were washed and stained for intracellular cytokines using BD cytofix/cytoperm kit (BD Pharmingen) according to the manufacturer’s instructions at the end of the 6-hour culture. Briefly, cells were first stained for surface markers, including CD4, CD8, and CD3, followed by intracellular staining with fluorophore-conjugated antibodies against IL-2, TNF-α, CD107a, and IFN-γ, or isotype-matched antibodies after fixation and permeabilization. In all stained samples, dead cells were excluded using LIVE/DEAD Fixable Aqua Dead Cell Staining Kit (Invitrogen, Carlsbad, CA).

### Antibodies

All antibodies used for flow cytometry analysis were purchased from BD Biosciences (San Jose, CA), Biolegend (San Diego, CA), Invitrogen (Carlsbad, CA) or eBioscience (Santa Clara, CA) and are listed in supplementary table 2.

### RNA-seq and data analysis

A total amount of 1 µg RNA per sample was used as input material for the RNA sample preparations. Sequencing libraries were generated using NEBNext® UltraTM RNA Library Prep Kit for Illumina® (NEB, USA) following manufacturer’s recommendations and index codes were added to attribute sequences to each sample. Briefly, mRNA was purified from total RNA using poly-T oligo-attached magnetic beads. Fragmentation was carried out using divalent cations under elevated temperature in NEBNext First Strand Synthesis Reaction Buffer (5X). First strand cDNA was synthesized using random hexamer primer and M-MuLV Reverse Transcriptase (RNase H-). Second strand cDNA synthesis was subsequently performed using DNA Polymerase I and RNase H. Remaining overhangs were converted into blunt ends via exonuclease/polymerase activities. After adenylation of 3’ ends of DNA fragments, NEBNext Adaptor with hairpin loop structure were ligated to prepare for hybridization. In order to select cDNA fragments of preferentially 150∼200 bp in length, the library fragments were purified with AMPure XP system (Beckman Coulter, Beverly, USA). Then 3 µl USER Enzyme (NEB, USA) was used with size-selected, adaptor-ligated cDNA at 37 °C for 15 min followed by 5 min at 95 °C before PCR. Then PCR was performed with Phusion High-Fidelity DNA polymerase, Universal PCR primers and Index (X) Primer. At last, PCR products were purified (AMPure XP system) and library quality was assessed on the Agilent Bioanalyzer 2100 system. The clustering of the index-coded samples was performed on a cBot Cluster Generation System using PE Cluster Kit cBot-HS (Illumina) according to the manufacturer’s instructions. After cluster generation, the library preparations were sequenced on an Illumina NextSeq 2000 platform and paired-end reads were generated. Raw data (reads) of FASTQ format were first processed to obtain clean reads by removing adapters, poly-N sequences and reads with low quality. The STAR aligner (v2.6.1) was then utilized to align the data to the mouse reference genome mm10, (Ensemble gene annotations). FeatureCounts (v1.5.0) was used to count the read numbers mapped of each gene and Reads Per Kilobase of exon model per Million mapped reads (RPKM) of each gene was calculated based on the length of the gene and reads count mapped to this gene ^27^. Differential expression analysis between two conditions/groups was performed using DESeq2 R package (v1.30.0). The resulting P values were adjusted using the Benjamini and Hochberg’s approach for controlling the False Discovery Rate (FDR). Genes with an adjusted P value < 0.05 found by DESeq2 were assigned as differentially expressed. Gene set enrichment analysis was performed with GSEA (v4.1.0) and hallmark, Gene Ontology, and KEGG gene sets obtained from Molecular Signatures Database (v7.4). We also conducted pathway enrichment analysis utilizing the GeneGo MetaCore+MetaDrug™ tool (Thompson Reuters, version 6.31 build 68930). Triplicate samples were initially processed. However, vehicle sample 3 for T cells and ACY241 sample 2 for TAMs were deemed as low quality and were excluded from further analysis. The RNA sequencing data have been deposited in the NCBI GEO public data repository (reference series accession GSEXXXX).

### Patient tumor 2-D cultures

Fresh resected NSCLC patient tumors were gently dissociated mechanically in RPMI 1640 media containing 10 U/mL Collagenase D (Sigma Aldrich) and 25 ug/mL DNase I grade II (Sigma Aldrich) followed by incubation at 37°C for 30 minutes. Dissociated tissue was further digested using GentleMACS (Miltenyi) after which single cells were passed over a 100 µm filter. Tumor cell suspensions were cultured in the presence of complete media supplemented with 10IU/ml of IL-2 and 10ng/ml of IL-15. ACY241 was added at a concentration of 100 nM while DMSO was added to control wells. After 4 days in culture, immunofluorescent staining was performed to determine the viability of tumor and immune cells by first staining with LIVE/DEAD dye followed by antibodies against EpCAM and CD45.

### ImageStream Analysis

Subcutaneous tumors were developed using B6 OVA 10103 F LT1 OVA-PGK cell line and tumor-bearing mice were treated with either vehicle or ACY241 for 2 weeks after which tumors were resected from mice and processed into cell suspensions. Samples then subsequently underwent staining with fluorophore-conjugated antibodies that included macrophage lineage markers as well as MHC class I and OVA OVA_257-264_-specific antibodies. Samples were acquired on the Amnis® ImageStream®X Mk II (Luminex) and analyzed with the IDEAS 6.2 software.

### Statistical analysis

Data were analyzed using mean ± standard error of the mean (SEM). Unpaired two-tailed Student *t* test was used for comparisons between two groups using GraphPad Prism software. *P* values < 0.05 were considered statistically significant (*); *P* values < 0.01 are marked **, and *P* values < 0.001 are marked ***.

## Results

### ACY241 treatment supports increased infiltration, activation, and effector profile of T cells in the lung tumors of a pre-clinical mouse model of NSCLC

Although previous report demonstrated the immunomodulatory properties of ricolinostat, an HDAC6 inhibitor in a murine model of NSCLC, its effects on tumor-associated immune cells did not translate to improved anti-tumor response as a monotherapy. This promising result nonetheless led us to evaluate ACY241, a structurally similar compound with higher potency for HDAC6 inhibition and favorable solubility and safety profile ^24 25^. We postulated that ACY241 administration in our pre-clinical mouse model of non-small cell lung cancer (NSCLC) will likely be associated with broader effects in the tumor microenvironment. In this regard, we first evaluated the nature and scope of ACY241 effects on tumor-associated immune cell subsets using the *Kras* mutant, *p53*-deficient (KP) genetically engineered mouse model of NSCLC in which spontaneous lung adenocarcinoma development is driven by activating *Kras* mutation and concurrent *p53* deficiency (denoted KP) ^26^.

Upon tumor establishment as confirmed by magnetic resonance imaging (MRI), mice were treated with ACY241 or vehicle as controls (Fig. 1A). Analysis of resected tumors after treatment cessation by multi-parameter flow cytometry revealed an increase in the proportions of CD4+Foxp3- and CD8+ effector T cells as well as NK cells in the tumors of ACY241-treated mice relative to the vehicle control group. In contrast, the proportions of CD4+CD25hiFoxp3+ regulatory T cells (Tregs) was diminished (Fig 1B). Myeloid cell subsets including tumor-associated macrophages (TAMs), myeloid-derived suppressor cells (MDSC), and dendritic cells (DC) were however not significantly altered (Fig. S1). Phenotypic assessment of CD8+ and CD4+Foxp3-T cells within the tumors of ACY241-treated mice showed that these cells exhibited enhanced activation status and proliferative profile evidenced by higher expression of CD69 and Ki67, respectively, compared to equivalent cells in the tumors of vehicle treated controls (Fig 1C, D, Fig. S2A). Furthermore, these CD8+ T cells also harbored increased central and effector memory phenotypic subsets (Fig. 1E and Fig. S2B).

**Figure 1.**
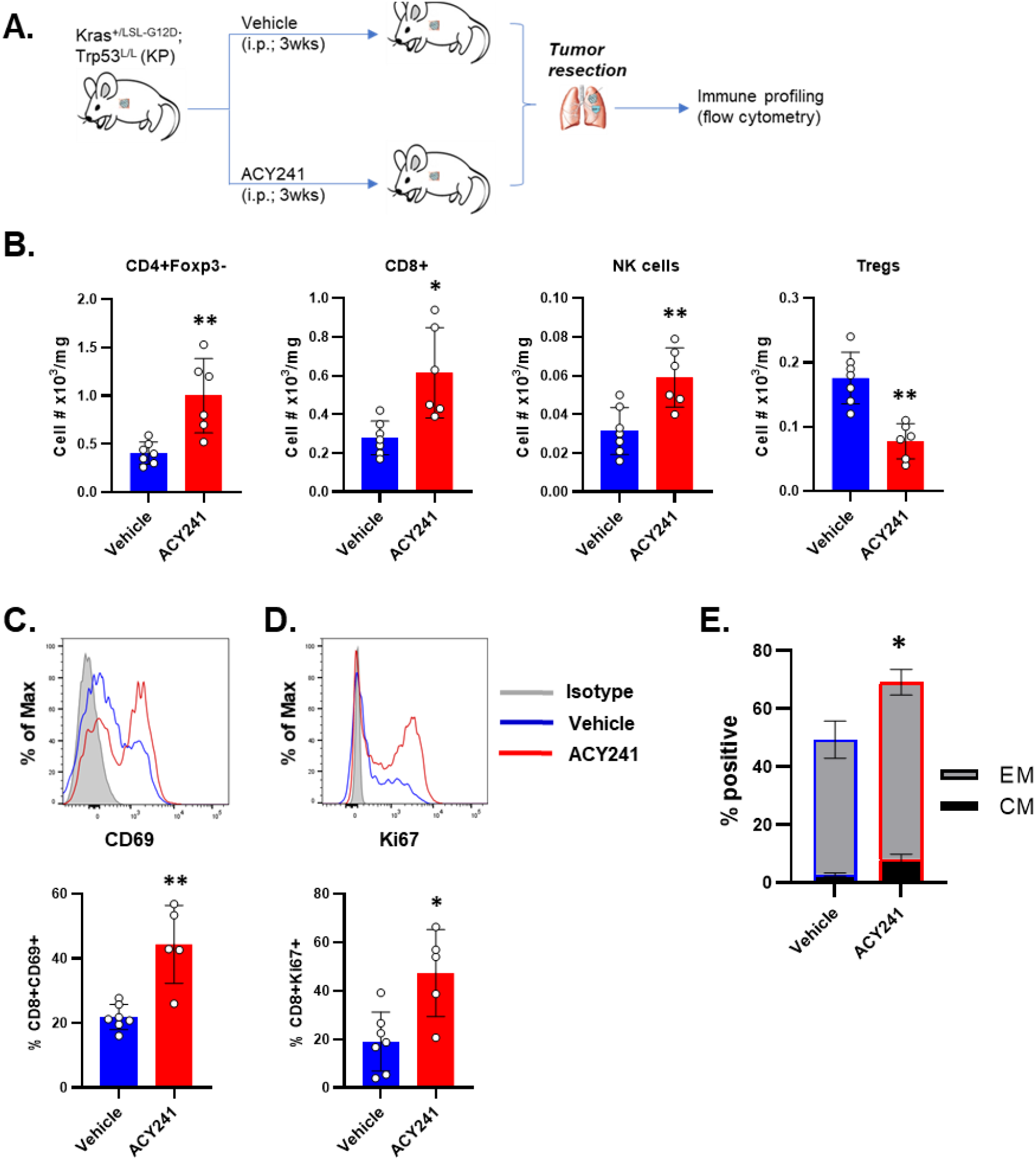
Increased infiltration and activation status of T cells in the tumors of ACY241-treated lung cancer-bearing mice. **(A)**. Schematics of treatment and analysis of lung tumor-bearing KP mice. Lung tumor formation was induced in *Kras*^+/LSL-G12D^; *Trp53*^L/L^ (KP) mice with intranasal administration of adeno-Cre. Mice with established tumors (150 – 200mm^3^) as confirmed by MRI were treated for three weeks with vehicle or ACY241 after which immune profiling was condcuted by flow cytometry to assess the numbers and phenotype of immune cell subsets. (B) Proportion of CD4+Foxp3-, CD8+ T cells, NK cells and Tregs (from left to right) within total viable cells per milligram of tumor. (C, D) Representative histograms top) and summary (bottom) of expression levels for (C) CD69 and (D) Ki67 on tumor-infiltrating CD8+ T cells. (E) Percent of CD8+ T cells with an effector memory (EM) or central memory (CM) phenotype within tumor infiltrating CD45+CD3+ cells as determined by CD62L and CD44 staining. Data in (E) are mean ±SEM of 5-7 mice per group. * indicates p-value < 0.05, ** p-value <0.01.

### ACY241 treatment is associated with reduced dysfunctional phenotype and enhanced effector capacity of NSCLC-infiltrating CD8+ T cells

Tumor-associated T cells exhibit a dysfunctional profile that is typified by sustained expression of an array of inhibitory proteins, a phenotype that is akin to a state of T cell exhaustion first described in chronic viral infections ^28^. Given existing reports demonstrating that HDAC6 inhibition can alter the expression of inhibitory checkpoint molecules in melanoma patient T cells ^29^, we evaluated the phenotype of lung adenocarcinoma-infiltrating T cells in our genetically engineered mouse model (GEMM) of NSCLC with particular focus on the expression pattern of inhibitory protein molecules as well as their effector function *ex vivo*. Our flow cytometric assessment revealed that in the presence of ACY241 treatment, lung tumor-infiltrating CD8+ T cells expressed lower levels of the inhibitory receptors BTLA, PD-1, and CTLA-4 relative to the CD8+ T cells in the tumors of vehicle-injected mice (Fig. 2A-C, Fig. S3A). As these inhibitory receptors contribute to T cell dysfunction ^30^, their diminished expression under ACY241 treatment raises the possibility that they may be less functionally impaired compared to equivalent cells in untreated control tumors. Indeed, when stimulated ex-vivo, a higher proportion of tumor isolated CD8+ (Fig. 2D, E) and CD4+ (Fig. S3B, C) T cells expressed CD107a, an indicator of degranulation, and granzyme B, a cytotoxic granule associated with effector function under ACY241 treatment. Collectively, these observations suggest that ACY241 promotes qualitative changes that is favorable to improved functionality of tumor-infiltrating T cells in NSCLC.

**Figure 2.**
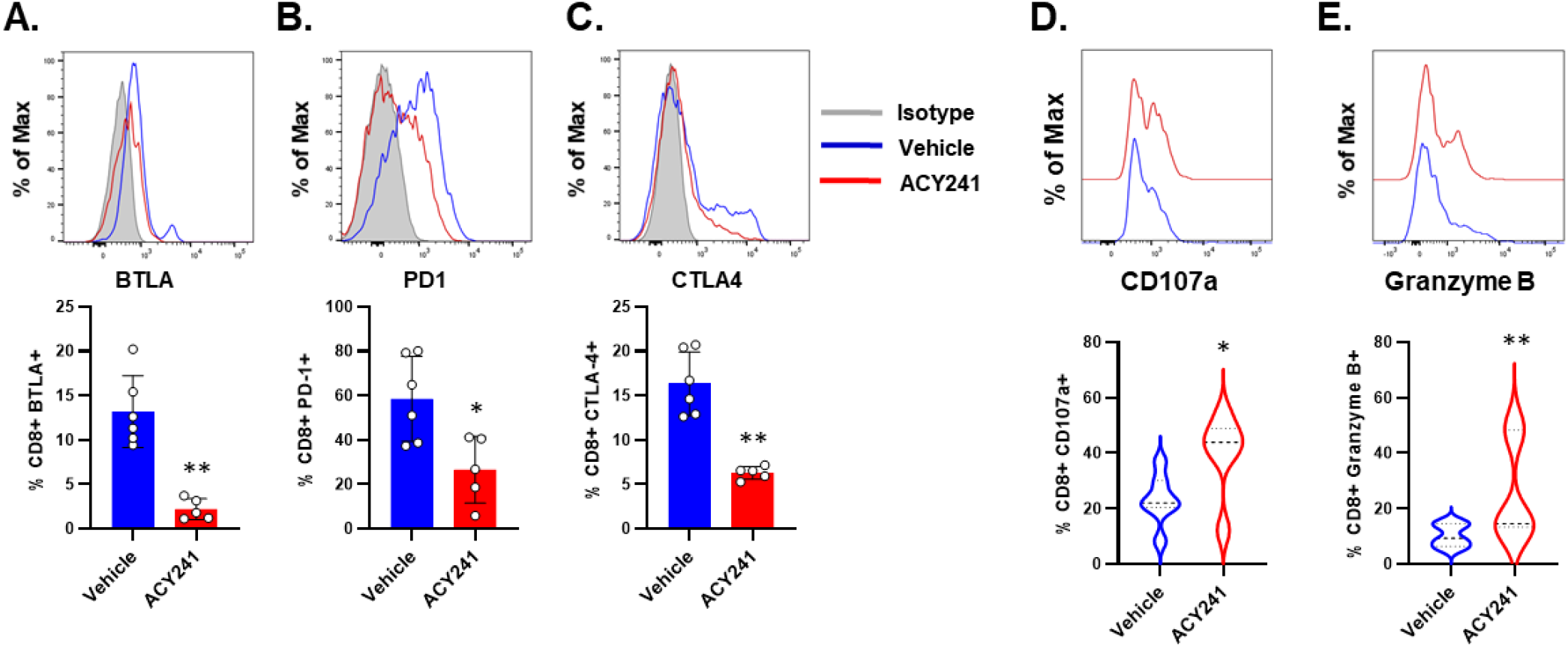
Reduced inhibitory receptor expression in tumor-infiltrating CD8+ T cells of ACY241-treated mice is accompanied by increased effector signature. Single cell suspensions generated from lung tumors of KP mice that were treated with vehicle or ACY241 were analysed by flow cytometry to determine the phenotype of tumor-associated CD45+CD3+ T cells. (A-C) Representative histograms (top) and summary (bottom) of expression levels for BTLA (A), PD-1 (B) and CTLA-4 (C) on tumor-infiltrating CD8+ T cell subset. T cells isolated from lung tumors of mice treated as indicated were stimulated for 6 hours with Cell stimulation cocktail plus Golgi inhibitor for subsequent intracellular cytokine staining. Representative histogram (top) and summary (bottom) for the expression of CD107a (D) or Granzyme B (E) on gated CD8+ T cells. Data in D and E (bottom) are mean ±SEM of 5-6 mice per group. * indicates p-value < 0.05, ** p-value <0.01.

### ACY241 treatment facilitates changes in tumor-associated macrophages that are conducive to enhanced antigenic stimulation of T cells

Besides the T cell compartment, we also interrogated the phenotype of myeloid cells, specifically tumor-associated macrophages (TAMs; CD11b+CD11c-Gr1-). First, we evaluated the expression of MHC, as well as co-stimulatory and co-inhibitory molecules in the presence or absence of ACY241 treatment. Our analysis showed that both MHC class I and II molecules were substantially increased in tumors of ACY241-treated mice relative to vehicle controls (Fig. 3 A, B) suggesting that ACY241 may facilitate improved presentation of tumor antigens to T cells in the tumor bed. Consistent with this idea, we found that TAMs in the tumors of ACY241-treated mice displayed higher expression of CD8+ T cell-specific OVA epitope (OVA_256-264_) in association with MHC class I (H-2Kb) in B6 mice bearing OVA-expressing tumors (Fig. 3C, D, Fig. S4A, B). We also observed a higher expression of this OVA antigenic peptide on the tumor cells in the ACY241 treatment condition (Fig. S4C). In additional phenotypic analysis, we found that ACY241-exposed TAMs displayed increased levels of co-stimulatory molecules CD80, CD86, and CD40 while expressing reduced levels of the inhibitory ligands PD-L1 and PD-L2 (Fig. 3E-I). These findings raise the possibility that ACY241 may tip the balance between activating and inhibitory signal availability to T cells in favor of the former in NSCLC.

**Figure 3.**
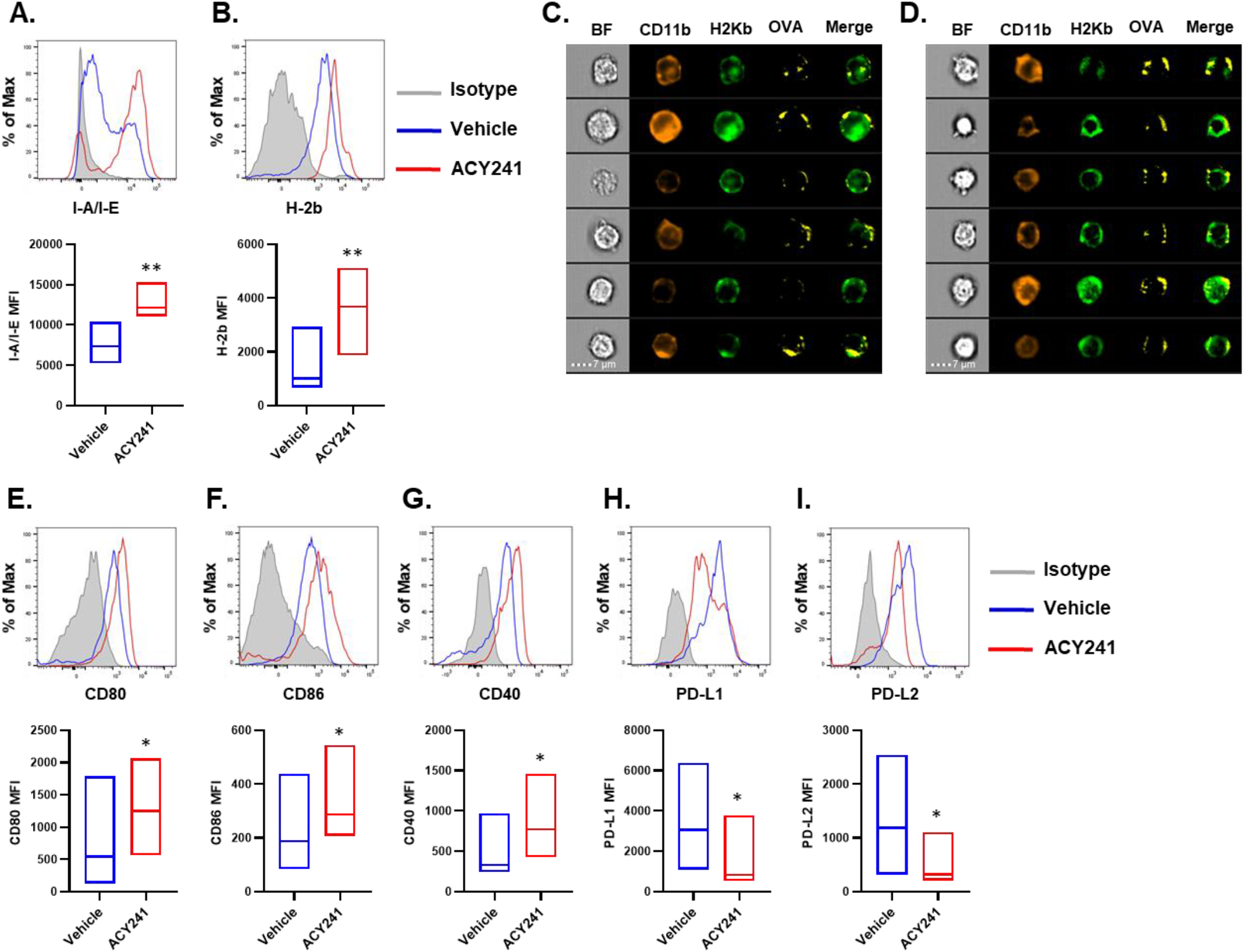
ACY241 potentiates phenotypic shift favoring upregulated expression of co-stimulatory molecules over inhibitory ligands in tumor-associated macrophages as well as tumor-associated antigens. The phenotype of tumor-associated macrophages (CD11b+CD11c-Gr-1-) in lung tumors of vehicle and ACY241-treated KP mice was evaluated by flow cytometry and imaging flowcytometry (Imagestream). Representative histograms (top) and summary (bottom) for the expression of (A) I-A/I-E (MHC class II), (B) H-2b (MHC class I). B6 mice were implanted with KP tumor cell line that expresses both CD4- and CD8-specific OVA epitopes. Mice were treated for 2 weeks with either vehicle or ACY241 after which tumor single cell suspensions were analyzed by imagestream. (C, D) Representative images for the expression of MHC class I H-2Kb and OVA_257-264_ on CD11b+CD11c-macrophages in the tumors of vehicle (C) versus ACY241 (D) treated mice. (E-G) Representative histograms (top) and summary (bottom) for the expression of (E) CD80, (F) CD86, (G) CD40, (H) PD-L1, and (I) PD-L2 on TAMs in vehicle or ACY241-treated KP mice as determined from median fluorescent intensity (MFI). Data are representative or are mean ±SEM of 5-7 mice per group. * indicates p-value < 0.05, ** p-value <0.01, *** p-value <0.001.

### ACY241 alters the transcriptional landscape of tumor-associated T cells and macrophages to reveal distinct genomic signatures accompanying its immunomodulatory effect

Although HDAC6 activity is often described in association with post-translational modification of its target proteins consistent with its largely cytoplasmic localization ^31-33^, increasing evidence support the idea that it likely has under-appreciated transcriptional regulatory activity ^34-36^. To gain some insight into the mechanism of ACY241 action in the context of transcriptional regulation that likely accompanies its immunomodulatory effects, we performed RNA-sequencing of T cells and TAMs that were sorted from the tumors of vehicle versus ACY241-treated KP mice. Our analysis reveals that only 213 genes were upregulated in ACY241-exposed tumor-T cells whereas over 4245 genes were downregulated (Log2 fold change greater than 1.5; SS Table 1). There was generally no significant effect on T cell-associated canonical genes that are often involved in T cell signaling, activation, differentiation and/or function except for *IL33* and *Nfatc2ip* (Fig. 4A, Fig. S5A). However, genes involved in apoptosis including *Fasl* and *Bax* were downregulated in ACY241-treated mice suggesting ACY241’s potential for enhancement of T cell survival. Notably, several genes encoding inhibitory receptors or molecules involved in attenuation of T cell optimal function such as *ctla4, vsir, tigit, lgals9, and Arg1* were also downregulated in tumor-infiltrating T cells exposed to ACY241 (Fig. 4A, Fig. S5A). Gene set enrichment analysis (GSEA) further revealed that genes involved in the PI3K/AKT/MTOR signaling pathway and TGF-Beta signaling were negatively enriched i.e. downregulated in tumor-T cells upon ACY241 treatment (Fig. 4B, C).

**Figure 4.**
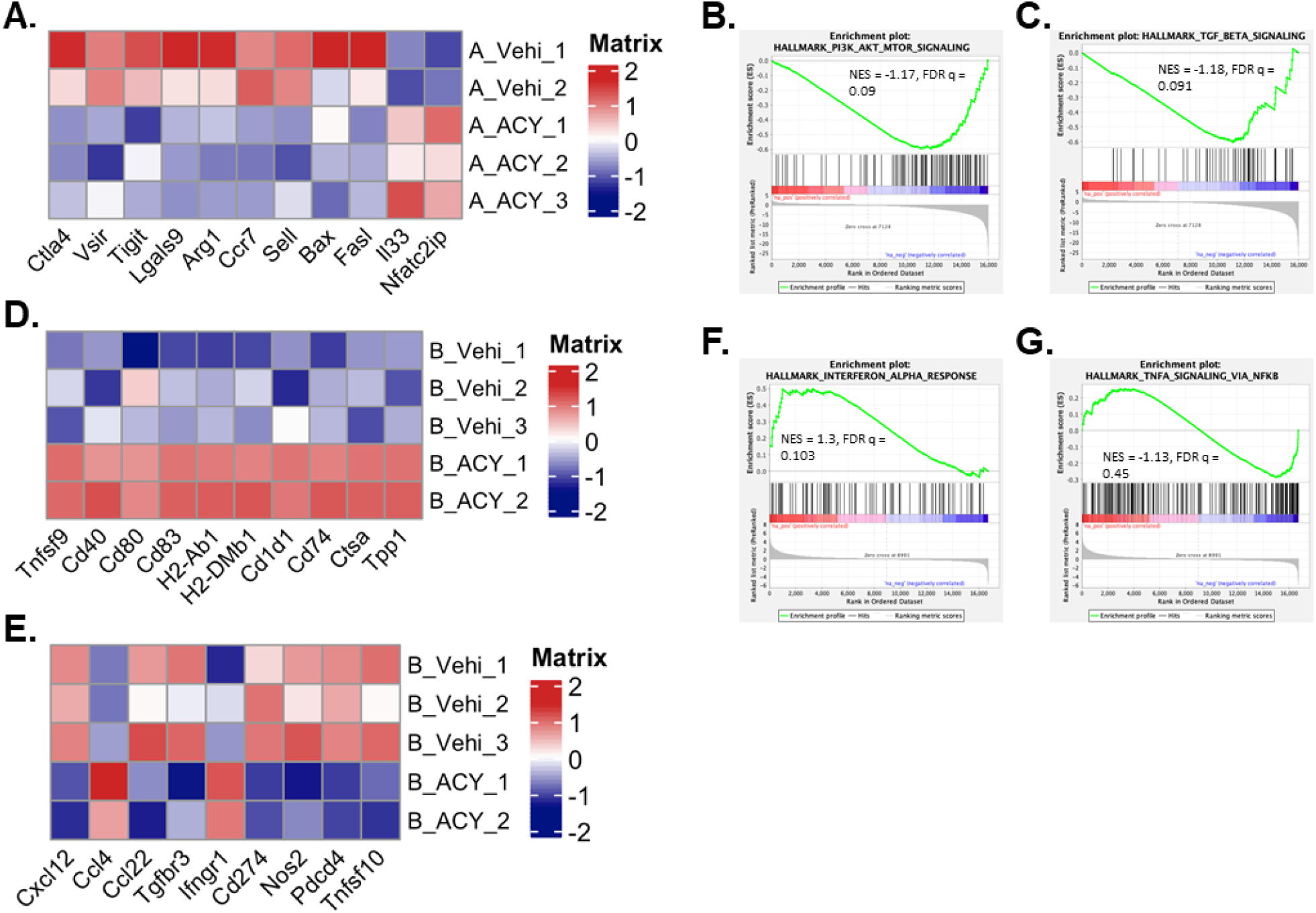
Transcriptomic analyses reveals genomic features that underlie the immunomodulatory activity of ACY241. CD45+CD3+Foxp3-T cells and CD45+CD11b+CD11c-Gr-1- tumor-associated macrophages (TAMs) were concurrently sorted from lung tumors of vehicle and ACY241-treated KP mice for bulk RNA sequencing on the NextSeq500 Illumina platform. Data was analyzed by the R package Seurat analysis pipeline. (A) Heat map for immune-related differentially expressed genes (≥ log2 fold change at padj value 0.05) in tumor-T cells from ACY241-treated mice relative to vehicle controls. Gene set enrichment analysis (GSEA) was performed to identify pathways highly enriched in tumor-T cells from ACY241-treated mice over vehicle. GSEA plot for (B) PI3K/AKT/MTOR and (C) TGF-beta signaling pathways. (D, E) Heat maps for immune-related differentially expressed genes (≥ log2 fold change at padj value 0.05) in TAMs from ACY241-treated mice relative to vehicle controls. (D) Expression pattern for indicated co-stimulatory, MHC, and related gene transcripts. (E) Expression pattern for indicated chemokines, chemokine receptor transcripts as well as indicated transcripts involved in attenuating T cell priming. (F, G) GSEA plots for (F) Interferon alpha response and (G) TNF-alpha signaling via NF-κB as determined by enrichment scores for gene sets in indicated pathways

Strikingly, in TAMs, 1,976 and 3,679 genes were up- or down-regulated, respectively after ACY241 treatment relative to vehicle demonstrating a more profound effect of ACY241 on TAMs compared to T cells (SS. Table 1, Fig. S5B). Among the most upregulated genes are those encoding costimulatory molecules Tnfrsf9, Cd40, Cd80, Cd83; MHC proteins H2-Ab1, H2-DMb1, as well as related proteins Cd1d1, Cd74, Ctsa, and Tpp1 (Fig. 4D). A number of chemokine genes such as *Cxcl12, Ccl4, and Ccl22* were also differentially expressed (Fig. 4 E). Furthermore, among the most prominently downregulated gene transcripts are genes whose protein derivatives play anti-inflammatory roles or function to attenuate T cell function. These include *Tgfbr3, Cd274, Nos, Pdcd4, and Tnfsf10* (Fig. 4 E, Fig. S5B). Further assessment by GSEA identified enrichment of genes that control Interferon alpha response and TNF-alpha signaling via NF-κB as well as JAK/STAT signaling pathway (Fig. 4F, G, Fig. S5C).

### Oxaliplatin synergizes with ACY241 to support enhanced anti-tumor response and tumor-associated T cell effector function

The chemotherapy drug oxaliplatin has been reported to promote immunogenic cell death of tumor cells ^37-39^. We reasoned that tumor-associated antigens arising from oxaliplatin-supported tumor cell death is likely to be presented in the context of enhanced MHC expression and costimulatory signals if partnered with ACY241 and this may support enhanced anti-tumor T cell response. To this end, we treated lung tumor-bearing KP mice 3x/week with ACY241 and/or Oxaliplatin. Control mice received vehicle. Survival of mice treated with ACY241 alone mirrored those treated with oxaliplatin although the former was slightly more efficacious in prolonging survival compared to vehicle-treated mice (MST 48 vs 41 vs 20). Administration of both agents concurrently achieved the most therapeutic outcome as mice treated with this combination survived much longer than those treated with either agent alone (MST 69) with some mice remaining alive past 80 days suggesting a synergistic effect from both drugs (Fig. 5A). Consistent with these observations, tumor weight as measured after 6 weeks of treatment was significantly reduced in ACY241 or Oxaliplatin-treated mice relative to vehicle control, and this was further reduced in the presence of the two drugs. (Fig. 5B). Surprisingly, we found that Gemcitabine, another chemotherapy drug that we evaluated with the rationale that it may synergize with ACY241 by lowering myeloid-derived suppressor cell (MDSC) numbers ^40^ did not promote any therapeutic benefit beyond what was associated with ACY241 alone. (Fig. S6). For mice monitored until clinical endpoints in the ACY241/oxaliplatin treatment cohorts, we conducted immune profiling of the resected lung tumors in individual mice in each group. We found significantly increased CD8+ T cells:Treg ratio in the tumors of ACY241+Oxaliplatin-treated mice which can be largely attributed to AC241 effect (Fig. 5C). Importantly, the capacity of these tumor-associated CD8+ T cells to produce IL-2, TNF-α, and IFN-γ upon ex-vivo stimulation was increased in ACY241 or oxaliplatin-treated mice compared to vehicle. This enhanced functional capability was further amplified upon combination of the two drugs. (Fig. 5D-F and G-I). Similar pattern was noted for the CD4+ T cells (Fig. S7). Thus, ACY241 cooperated with oxaliplatin to enhance polyfunctionality of tumor-associated T cells and promote a durable anti-tumor response in our pre-clinical model of NSCLC. To determine if this particular drug combination has deleterious effects on tumor-associated immune cells which are important mediators of durable anti-tumor responses ^41^, we evaluated the viability of tumor and immune cells in a 2-D culture of freshly resected NSCLC patient tumor in the presence or absence of ACY241 and/or oxaliplatin. While reduced tumor cell viability followed a decreasing trend from ACY241 to oxaliplatin to the combination, immune cells were largely unaffected in the presence of one or both drugs in this short-term culture (Fig. S8) suggesting that the combination of ACY241 and oxaliplatin does not promote a significant toxic effect on tumor-associated immune cells.

**Figure 5.**
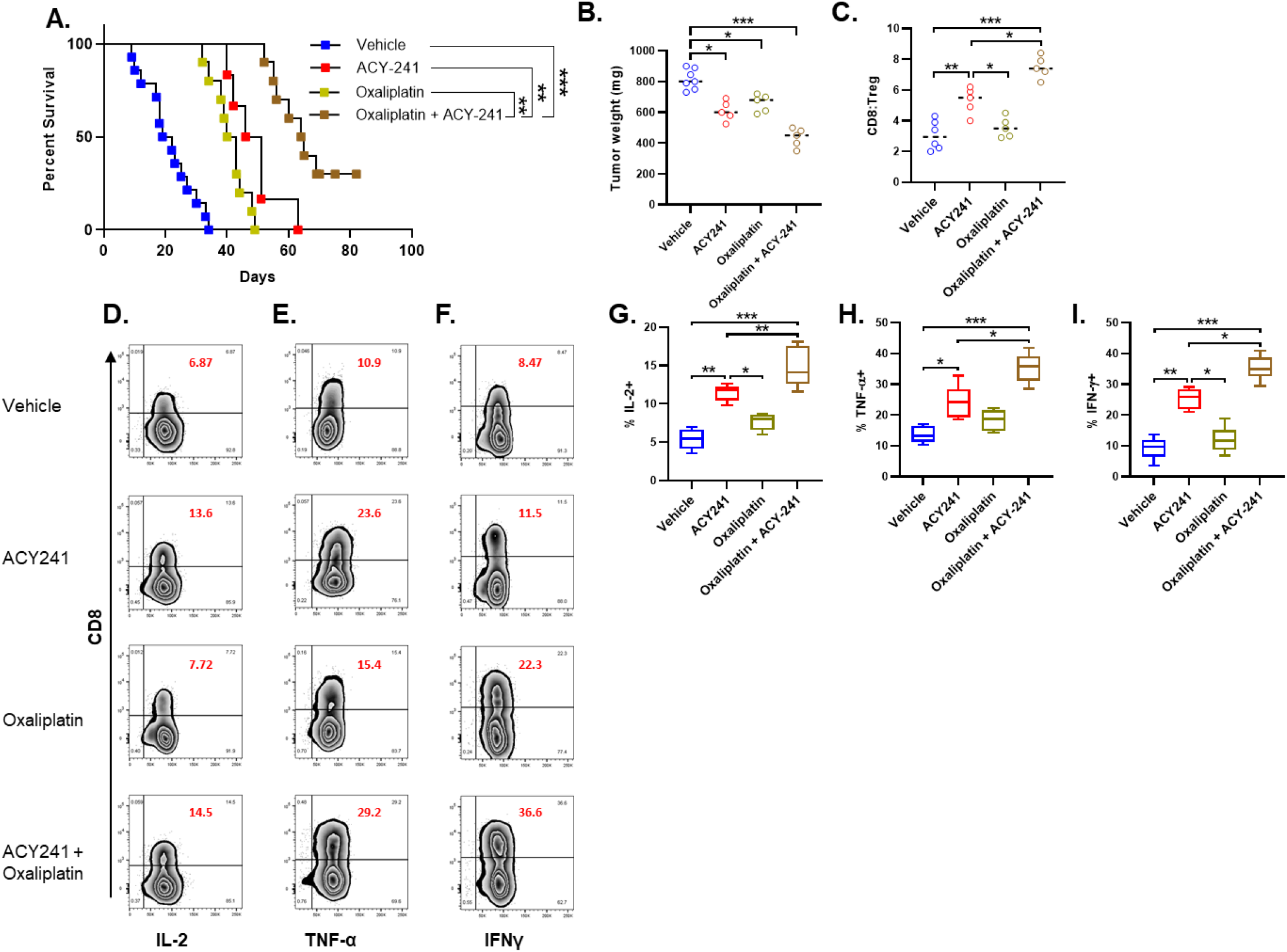
ACY241 coupled with Oxaliplatin significantly improves the survival of tumor-bearing mice. KP mice with established tumors (150 – 200mm^3^) received intraperitoneal injections of ACY241 (3X/week) and/or Oxaliplatin (1x/week) for 6 consecutive weeks. Mice receiving vehicle served as controls. One cohort was analyzed right after treatment cessation while a second cohort was monitored until clinical endpoints. (A) Kinetics of survival for tumor-bearing mice that were treated as indicated and monitored long-term. (B) Weight of tumor nodules in the lungs of treated mice analysed right after 6-week treatment. Immune profiling was conducted by flow cytometry using tumor single cell suspensions to determine proportion of T cell subsets. (C) CD8+ T cells to Tregs (CD4+CD25hiFoxp3+) ratio in tumors of mice in each treatment group. Tumor-infiltrating CD45+CD3+ T cells were isolated from tumor cell suspensions and equivalent numbers stimulated for 6 hours with Cell stimulation cocktail plus Golgi inhibitor for subsequent intracellular staining. Representative Zebra plot (D-F) or summary (G-I) for proportion of gated CD8+ T cells secreting IL-2 (D, G), TNF-α (E, H), and IFNγ (F, I) under each treatment condition as indicated. Data are representative (D-F) or are mean ± SEM (A, G-I) of 5-7 mice per group. * indicates p-value < 0.05, ** p-value <0.01, *** p-value <0.001.

## Discussion

In hematologic cancers, both pan and selective HDAC inhibitors show promising therapeutic results ^42 43^. However, solid cancers have not met with the same level of success although emerging data is encouraging ^6 16 22-25 44^. In the present study, we evaluated ACY241, an HDAC6-selective inhibitor in a pre-clinical mouse model of NSCLC to determine its therapeutic potential in this disease by leveraging its identified immunomodulatory effects in new combinatorial drug regimens.

Our immunogenomic studies suggest that ACY241 has a broad effect on NSCLC-associated macrophages and T cells. The sheer number of differentially regulated genes in the tumor-associated macrophages and T cells support this notion. Indeed, several genes involved in signaling, apoptosis, MHC assembly machinery, antigen presentation, co-inhibition and co-stimulation were differentially altered in these cells upon ACY241 treatment relative to vehicle controls. The upregulation of gene sets involved in the PI3K/AKT/MTOR pathway in tumor-associated T cells suggests that HDAC6 may play a critical role in regulating T cell function through this signaling pathway. On the other hand, the upregulation of IFN-α and TNF-α signaling pathway-associated gene sets in macrophages raises the possibility that AC241 may regulate macrophage activity in inflammatory responses in general including innate immune responses against pathogens such as viral infections.

While HDAC6 activity is primarily cytosolic ^31 32^, our study supports the growing body of evidence that it exhibits transcriptional regulatory activity and it likely interacts with other transcriptional factors, co-factors, and regulators to modulate transcriptional machinery associated with a number of genes ^34-36^. Indeed, HDAC6 interactions with Stat3 and the IL-10 gene promoter has been described ^45^.

Our findings are congruent with an expansive immunostimulatory activity of ACY241 beyond what has been previously appreciated. This includes its facilitation of increased T and NK cell numbers in the tumor microenvironment (TME). For the former, it remains a possibility that ACY241 promotes clonal expansion of tumor-reactive T cells. To our knowledge, no HDAC inhibitor has been reported to selectively expand antigen-specific or clonal populations of T cells in tumors. Nonetheless, we speculate that such outcome may arise through indirect effect whereby the composite of ACY241-associated increase in MHC expression, co-stimulatory molecules and decrease in inhibitory ligands in APCs lowers the threshold for activation and priming of effector T cells with varying degrees of affinities, facilitating their expansion in the tumor. This is plausible as tumor-associated T cells under ACY241 treatment exhibited increased proliferative profile compared to vehicle-treated group. Future studies focused on analysis of the T cell receptor repertoire of tumor-infiltrating T cells in the presence or absence of ACY241 treatment could shed light on this possibility.

Tumor antigen vaccinations have been explored as a modality for immunotherapy of cancer ^46 47^. While tumor-associated antigens that are immunogenic have not been fully elucidated in lung cancer, our findings that ACY241 promotes broad phenotypic changes in both tumor-infiltrating T cells and TAMs that in principle should promote enhanced antigen presentation and co-activation of T cells supports its suitability in tumor vaccination settings. Given the observation that the expression of tumor-expressed OVA antigenic epitopes associated with MHC molecules was not only higher on tumor macrophages upon ACY241 treatment but was also upregulated on tumor cells, it is likely that ACY241 promotes increased immunogenicity of lung adenocarcinoma cells. This is consistent with the report by others in which the HDAC inhibitor LBH589 promoted upregulation of tumor-associated antigen-encoding transcripts in a melanoma model ^19^.

Although its therapeutic effect was modest as a single agent in our studies, we speculate that it is likely to be a suitable rationally selected partner drug in novel combinatorial drug regimens where the other agent precipitates increased availability of tumor antigens through immunogenic tumor cell death. Our results from its combination with oxaliplatin indeed is in alignment with this idea. Given the toxicity often associated with chemotherapy agents such as oxaliplatin ^48 49^, it was somewhat surprising to us that the combination of ACY241 and oxaliplatin was well tolerated in the treated mice with no observable evidence of severe toxicity. Thus, this regimen appears to show a relatively safe profile at least in this pre-clinical mouse model. In support of this inference, tumor-associated leukocytes did not show impaired viability in the presence of both agents *in vitro* unlike tumor cells. Thus, we opine that the combination of ACY241 and oxaliplatin is a promising and potentially tolerable regimen worth evaluating in the clinic for NSCLC therapy.

Our studies raise an important point; that HDAC6 likely has broad transcriptional activity in tumor-associated immune T cells and macrophages, and this may not necessarily be a tumor-specific phenomenon, but one that likely applies to the function of these cells in other inflammatory settings necessitating innate and adaptive immune responses. Of note is the regulation of a number of immune checkpoint genes which were downregulated in tumor-infiltrating T cells upon ACY241 treatment suggesting its transcriptional regulatory activity may extend beyond what we have previously appreciated. The notion that ACY241 could be utilized to potentially rescue “exhausted” T cells by downregulating their inhibitory molecular signature is intriguing in light of the observation that the functional capacity of tumor-T cells in mice treated with ACY241 improved significantly concomitant with reduced expression of exhaustion-implicated proteins. Beyond its overall therapeutic appeal in cancer, its utility to potentially reprogram partially exhausted T cells to a more functional state is a subject of our follow up studies.

Although ACY241 has been tested in clinical trials in combination with anti-PD-1 or paclitaxel (NCT02635061, NCT02551185) ^50^, it’s therapeutic potential in NSCLC is still being explored. Our observations support the notion that it has robust activity as a non-prototypic immunomodulatory agent. Thus, rational selection of partner drugs may cooperate, synergize, or amplify the positive immunostimulatory effects of ACY241 to a threshold that will support therapeutic outcomes. On this note, our studies suggest that its therapeutic potential will likely be maximized when partnered with agents that capitalize on what it appears to do efficiently – enhance effector cell infiltration, support of macrophage-mediated T cell priming and lowering molecular brakes on T cells.

## Supporting information

Cover letter to JITC

## Abbreviations

cDNA: Complementary DNA
DC: Dendritic cell
FcR: Fc receptor
FDR: False discovery rate
GEMM: Genetically engineered mouse model
HDAC: Histone deacetylase
MDSC: Myeloid-derived suppressor cell
NK: Natural killer
NSCLC: Non-small cell lung cancer
OVA: Ovalbumin
TAM: Tumor-associated macrophage
TME: Tumor microenvironment
Treg: Regulatory T-cells

## Acknowledgements

We thank the Small animal imaging core facility for their contribution to MRI imaging of tumor-bearing mice, Olya Stringfield for her assistance with MRI image analysis, Neel Chaudhary for assistance with ImageStream analysis, the flow cytometry core for their support of sample acquisition, and the bioinformatics core for RNA-sequencing data analysis. Special thanks to Dr. Tyler Jacks for the generous gift of the B6 OVA 10103 F LT1 OVA-PGK tumor cell line.

## Funding

This work was supported by NCI 1K22CA222669-01 (to DA), American lung Association award ALA69-20210-02-01 (to D.A) and in part by the Flow Cytometry Core Facility at the Moffitt Cancer Center, an NCI designated Comprehensive Cancer Center (P30-CA076292).

## Availability of data and materials

The data generated and/or analyzed in this study are included in this published article (and its supplementary information files). The RNA-Seq data generated and/or reported in this study will be available in the Gene Expression Omnibus (GEO) repository upon acceptance for publication.

## Author contributions

AB, AS, SB, WD, DA: conducted experiments

AB, AS, LC, DA: analyzed data.

AB, DA: interpreted data, edited the manuscript.

AB, LC, DA: discussed data and manuscript.

DA: designed research studies, discussed data sets, wrote the paper, supervised, and supported the research.

## Ethics Declarations

### Ethics approval

All breeding and treatment studies were performed with the approval of Moffitt Cancer Center Animal Care and Use Committee. All animal work was conducted in accordance with ARRIVE guidelines and in accordance with institutional guidelines for animal welfare.

### Consent for publication

Not applicable

### Competing interests

All authors declare no competing interests.

## Notes

### Competing Interest Statement

The authors have declared no competing interest.

## References

1. Garon EB, Rizvi NA, Hui R, et al. Pembrolizumab for the treatment of non-small-cell lung cancer. N Engl J Med 2015;372(21):2018–28. doi: 10.1056/NEJMoa1501824 [published Online First: 2015/04/22]

2. Aguilar EJ, Ricciuti B, Gainor JF, et al. Outcomes to first-line pembrolizumab in patients with non-small-cell lung cancer and very high PD-L1 expression. Ann Oncol 2019;30(10):1653–59. doi: 10.1093/annonc/mdz288 [published Online First: 2019/08/23]

3. Carbone DP, Reck M, Paz-Ares L, et al. First-Line Nivolumab in Stage IV or Recurrent Non-Small-Cell Lung Cancer. N Engl J Med 2017;376(25):2415–26. doi: 10.1056/NEJMoa1613493 [published Online First: 2017/06/22]

4. Hellmann MD, Ciuleanu TE, Pluzanski A, et al. Nivolumab plus Ipilimumab in Lung Cancer with a High Tumor Mutational Burden. N Engl J Med 2018;378(22):2093–104. doi: 10.1056/NEJMoa1801946 [published Online First: 2018/04/17]

5. Hellmann MD, Paz-Ares L, Bernabe Caro R, et al. Nivolumab plus Ipilimumab in Advanced Non-Small-Cell Lung Cancer. N Engl J Med 2019;381(21):2020–31. doi: 10.1056/NEJMoa1910231 [published Online First: 2019/09/29]

6. Lane AA, Chabner BA. Histone deacetylase inhibitors in cancer therapy. J Clin Oncol 2009;27(32):5459–68. doi: 10.1200/JCO.2009.22.1291 [published Online First: 2009/10/15]

7. Li Y, Seto E. HDACs and HDAC Inhibitors in Cancer Development and Therapy. Cold Spring Harb Perspect Med 2016;6(10) doi: 10.1101/cshperspect.a026831 [published Online First: 2016/09/08]

8. Panobinostat Approved for Multiple Myeloma. Cancer Discovery 2015(10.1158/2159-8290.CD-NB2015-040) doi: 10.1158/2159-8290.CD-NB2015-040

9. Nooka AK, Kastritis E, Dimopoulos MA, et al. Treatment options for relapsed and refractory multiple myeloma. Blood 2015;125(20):3085–99. doi: 10.1182/blood-2014-11-568923 [published Online First: 2015/04/04]

10. Mann BS, Johnson JR, Cohen MH, et al. FDA approval summary: vorinostat for treatment of advanced primary cutaneous T-cell lymphoma. Oncologist 2007;12(10):1247–52. doi: 10.1634/theoncologist.12-10-1247 [published Online First: 2007/10/27]

11. Qiu T, Zhou L, Zhu W, et al. Effects of treatment with histone deacetylase inhibitors in solid tumors: a review based on 30 clinical trials. Future Oncol 2013;9(2):255–69. doi: 10.2217/fon.12.173 [published Online First: 2013/02/19]

12. Anne M, Sammartino D, Barginear MF, et al. Profile of panobinostat and its potential for treatment in solid tumors: an update. Onco Targets Ther 2013;6:1613–24. doi: 10.2147/OTT.S30773 [published Online First: 2013/11/23]

13. Ceccacci E, Minucci S. Inhibition of histone deacetylases in cancer therapy: lessons from leukaemia. Br J Cancer 2016;114(6):605–11. doi: 10.1038/bjc.2016.36 [published Online First: 2016/02/26]

14. Imai Y, Maru Y, Tanaka J. Action mechanisms of histone deacetylase inhibitors in the treatment of hematological malignancies. Cancer Sci 2016;107(11):1543–49. doi: 10.1111/cas.13062 [published Online First: 2016/08/25]

15. Li G, Tian Y, Zhu WG. The Roles of Histone Deacetylases and Their Inhibitors in Cancer Therapy. Front Cell Dev Biol 2020;8:576946. doi: 10.3389/fcell.2020.576946 [published Online First: 2020/10/30]

16. Woods DM, Sodre AL, Villagra A, et al. HDAC Inhibition Upregulates PD-1 Ligands in Melanoma and Augments Immunotherapy with PD-1 Blockade. Cancer Immunol Res 2015;3(12):1375–85. doi: 10.1158/2326-6066.CIR-15-0077-T [published Online First: 2015/08/25]

17. West AC, Christiansen AJ, Smyth MJ, et al. The combination of histone deacetylase inhibitors with immune-stimulating antibodies has potent anti-cancer effects. Oncoimmunology 2012;1(3):377–79. doi: 10.4161/onci.18804 [published Online First: 2012/06/28]

18. Khan AN, Gregorie CJ, Tomasi TB. Histone deacetylase inhibitors induce TAP, LMP, Tapasin genes and MHC class I antigen presentation by melanoma cells. Cancer Immunol Immunother 2008;57(5):647–54. doi: 10.1007/s00262-007-0402-4 [published Online First: 2007/11/30]

19. Woods DM, Woan K, Cheng F, et al. The antimelanoma activity of the histone deacetylase inhibitor panobinostat (LBH589) is mediated by direct tumor cytotoxicity and increased tumor immunogenicity. Melanoma Res 2013;23(5):341–8. doi: 10.1097/CMR.0b013e328364c0ed [published Online First: 2013/08/22]

20. Armeanu S, Bitzer M, Lauer UM, et al. Natural killer cell-mediated lysis of hepatoma cells via specific induction of NKG2D ligands by the histone deacetylase inhibitor sodium valproate. Cancer Res 2005;65(14):6321–9. doi: 10.1158/0008-5472.CAN-04-4252 [published Online First: 2005/07/19]

21. Licciardi PV, Karagiannis TC. Regulation of immune responses by histone deacetylase inhibitors. ISRN Hematol 2012;2012:690901. doi: 10.5402/2012/690901 [published Online First: 2012/03/31]

22. Zheng H, Zhao W, Yan C, et al. HDAC Inhibitors Enhance T-Cell Chemokine Expression and Augment Response to PD-1 Immunotherapy in Lung Adenocarcinoma. Clin Cancer Res 2016;22(16):4119–32. doi: 10.1158/1078-0432.CCR-15-2584 [published Online First: 2016/03/12]

23. Bae J, Hideshima T, Tai YT, et al. Histone deacetylase (HDAC) inhibitor ACY241 enhances anti-tumor activities of antigen-specific central memory cytotoxic T lymphocytes against multiple myeloma and solid tumors. Leukemia 2018;32(9):1932–47. doi: 10.1038/s41375-018-0062-8 [published Online First: 2018/03/01]

24. Huang P, Almeciga-Pinto I, Jarpe M, et al. Selective HDAC inhibition by ACY-241 enhances the activity of paclitaxel in solid tumor models. Oncotarget 2017;8(2):2694–707. doi: 10.18632/oncotarget.13738 [published Online First: 2016/12/08]

25. North BJ, Almeciga-Pinto I, Tamang D, et al. Enhancement of pomalidomide anti-tumor response with ACY-241, a selective HDAC6 inhibitor. PLoS One 2017;12(3):e0173507. doi: 10.1371/journal.pone.0173507 [published Online First: 2017/03/07]

26. DuPage M, Dooley AL, Jacks T. Conditional mouse lung cancer models using adenoviral or lentiviral delivery of Cre recombinase. Nat Protoc 2009;4(7):1064–72. doi: 10.1038/nprot.2009.95 [published Online First: 2009/06/30]

27. Mortazavi A, Williams BA, McCue K, et al. Mapping and quantifying mammalian transcriptomes by RNA-Seq. Nat Methods 2008;5(7):621–8. doi: 10.1038/nmeth.1226 [published Online First: 2008/06/03]

28. Wherry EJ, Kurachi M. Molecular and cellular insights into T cell exhaustion. Nat Rev Immunol 2015;15(8):486–99. doi: 10.1038/nri3862 [published Online First: 2015/07/25]

29. Laino AS, Betts BC, Veerapathran A, et al. HDAC6 selective inhibition of melanoma patient T-cells augments anti-tumor characteristics. J Immunother Cancer 2019;7(1):33. doi: 10.1186/s40425-019-0517-0 [published Online First: 2019/02/08]

30. Thommen DS, Schumacher TN. T Cell Dysfunction in Cancer. Cancer Cell 2018;33(4):547–62. doi: 10.1016/j.ccell.2018.03.012 [published Online First: 2018/04/11]

31. Verdel A, Curtet S, Brocard MP, et al. Active maintenance of mHDA2/mHDAC6 histone-deacetylase in the cytoplasm. Curr Biol 2000;10(12):747–9. doi: 10.1016/s0960-9822(00)00542-x [published Online First: 2000/06/30]

32. Bertos NR, Gilquin B, Chan GK, et al. Role of the tetradecapeptide repeat domain of human histone deacetylase 6 in cytoplasmic retention. J Biol Chem 2004;279(46):48246–54. doi: 10.1074/jbc.M408583200 [published Online First: 2004/09/07]

33. Boyault C, Sadoul K, Pabion M, et al. HDAC6, at the crossroads between cytoskeleton and cell signaling by acetylation and ubiquitination. Oncogene 2007;26(37):5468–76. doi: 10.1038/sj.onc.1210614 [published Online First: 2007/08/19]

34. Westendorf JJ, Zaidi SK, Cascino JE, et al. Runx2 (Cbfa1, AML-3) interacts with histone deacetylase 6 and represses the p21(CIP1/WAF1) promoter. Mol Cell Biol 2002;22(22):7982–92. doi: 10.1128/MCB.22.22.7982-7992.2002 [published Online First: 2002/10/23]

35. Zhang W, Kone BC. NF-kappaB inhibits transcription of the H(+)-K(+)-ATPase alpha(2)-subunit gene: role of histone deacetylases. Am J Physiol Renal Physiol 2002;283(5):F904–11. doi: 10.1152/ajprenal.00156.2002 [published Online First: 2002/10/10]

36. Fernandes I, Bastien Y, Wai T, et al. Ligand-dependent nuclear receptor corepressor LCoR functions by histone deacetylase-dependent and -independent mechanisms. Mol Cell 2003;11(1):139–50. doi: 10.1016/s1097-2765(03)00014-5 [published Online First: 2003/01/22]

37. Wang YJ, Fletcher R, Yu J, et al. Immunogenic effects of chemotherapy-induced tumor cell death. Genes Dis 2018;5(3):194–203. doi: 10.1016/j.gendis.2018.05.003 [published Online First: 2018/10/16]

38. Tesniere A, Schlemmer F, Boige V, et al. Immunogenic death of colon cancer cells treated with oxaliplatin. Oncogene 2010;29(4):482–91. doi: 10.1038/onc.2009.356 [published Online First: 2009/11/03]

39. Shalapour S, Font-Burgada J, Di Caro G, et al. Immunosuppressive plasma cells impede T-cell-dependent immunogenic chemotherapy. Nature 2015;521(7550):94–8. doi: 10.1038/nature14395 [published Online First: 2015/04/30]

40. Sawant A, Schafer CC, Jin TH, et al. Enhancement of antitumor immunity in lung cancer by targeting myeloid-derived suppressor cell pathways. Cancer Res 2013;73(22):6609–20. doi: 10.1158/0008-5472.CAN-13-0987 [published Online First: 2013/10/03]

41. Fridman WH, Zitvogel L, Sautes-Fridman C, et al. The immune contexture in cancer prognosis and treatment. Nat Rev Clin Oncol 2017;14(12):717–34. doi: 10.1038/nrclinonc.2017.101 [published Online First: 2017/07/26]

42. Chen IC, Sethy B, Liou JP. Recent Update of HDAC Inhibitors in Lymphoma. Front Cell Dev Biol 2020;8:576391. doi: 10.3389/fcell.2020.576391 [published Online First: 2020/10/06]

43. Pride DA, Summers AR. The emergence of specific HDAC inhibitors and their clinical efficacy in the treatment of hematologic malignancies and breast cancer. Int J Mol Biol Open Access 2018;3(5):7. doi: 10.15406/ijmboa.2018.03.00078 [published Online First: September 26]

44. West AC, Johnstone RW. New and emerging HDAC inhibitors for cancer treatment. J Clin Invest 2014;124(1):30–9. doi: 10.1172/JCI69738 [published Online First: 2014/01/03]

45. Cheng F, Lienlaf M, Wang HW, et al. A novel role for histone deacetylase 6 in the regulation of the tolerogenic STAT3/IL-10 pathway in APCs. J Immunol 2014;193(6):2850–62. doi: 10.4049/jimmunol.1302778 [published Online First: 2014/08/12]

46. Peng M, Mo Y, Wang Y, et al. Neoantigen vaccine: an emerging tumor immunotherapy. Mol Cancer 2019;18(1):128. doi: 10.1186/s12943-019-1055-6 [published Online First: 2019/08/25]

47. Gupta RG, Li F, Roszik J, et al. Exploiting Tumor Neoantigens to Target Cancer Evolution: Current Challenges and Promising Therapeutic Approaches. Cancer Discov 2021;11(5):1024–39. doi: 10.1158/2159-8290.CD-20-1575 [published Online First: 2021/03/17]

48. Hoff PM, Saad ED, Costa F, et al. Literature review and practical aspects on the management of oxaliplatin-associated toxicity. Clin Colorectal Cancer 2012;11(2):93–100. doi: 10.1016/j.clcc.2011.10.004 [published Online First: 2011/12/14]

49. Mitchell EP. Gastrointestinal toxicity of chemotherapeutic agents. Semin Oncol 2006;33(1):106–20. doi: 10.1053/j.seminoncol.2005.12.001 [published Online First: 2006/02/14]

50. Awad MM, Le Bruchec Y, Lu B, et al. Selective Histone Deacetylase Inhibitor ACY-241 (Citarinostat) Plus Nivolumab in Advanced Non-Small Cell Lung Cancer: Results From a Phase Ib Study. Front Oncol 2021;11:696512. doi: 10.3389/fonc.2021.696512 [published Online First: 2021/09/24]

