## Supplemental Tables and Figures for "The immunomodulatory properties of the HDAC6 inhibitor ACY241 supports robust anti-tumor response in NSCLC when coupled with the chemotherapy drug Oxaliplatin"

**Supplementary Materials**

**Table S1.**

| | T-cell | TAM |
|---|---|---|
| Up-regulated genes* | 213 | 1976 |
| Down-regulated genes* | 4245 | 3679 |
| *Cutoff: adjust p-value < 0.05 and > 2 fold-change | | |

**Table S1. Differential expression of gene transcripts in ACY241-treated mice.** Bulk RNA-sequencing was performed on T cells and TAMs isolated from the tumors of mice treated with ACY241 or vehicle as controls. Number of upregulated or downregulated genes in tumor-infiltrating effector T cells and Tumor-associated macrophages (TAM) in ACY241-treated samples as determined based on Log2FC over vehicle and padj cut-off of 0.05.

**Table S2.**

| Fluorescent label | Antigen | Clone | Supplier |
|---|---|---|---|
| FITC | Gr-1 | RB6-8C5 | Biolegend |
| | CD107a | 1D4B | Biolegend |
| | H2 | M1/42 | Biolegend |
| Percp-Cy5.5 | CD8 | 53-6.7 | Biolegend |
| | PD-L2 | TY25 | Biolegend |
| | CD11c | N418 | Biolegend |
| APC | Nkp46 | 29AL4 | Biolegend |
| | BTLA | 6A6 | Biolegend |
| | GZB | QA16A02 | Biolegend |
| | PD-L1 | 10F.9G2 | Biolegend |
| AF700 | Foxp3 | FJK-16s | Invitrogen |
| | IA/IE | M5/114.15.2 | Biolegend |
| APC/Cy7 | CD11c | N418 | Biolegend |
| | CD44 | IM7 | Biolegend |
| BV-421 | F4/80 | BM8 | Biolegend |
| | Ki67 | 16A8 | Biolegend |
| | TNF-α | MP6-XT22 | Biolegend |
| BV605 | CD45 | 30-F11 | Biolegend |
| BV650 | CD4 | RM4-5 | Biolegend |
| | CD80 | 16-10A1 | Biolegend |
| BV711 | CD40 | 3/23 | BD Bioscience |
| BV785 | CD3 | 17A2 | Biolegend |
| BUV395 | CD25 | PC61 | BD Bioscience |
| | CD47 | miap301 | BD Bioscience |
| BUV737 | PD1 | J43 | BD Bioscience |
| | EpCAM | G8.8 | BD Bioscience |
| PE | CD19 | 6D5 | Biolegend |
| | CTLA4 | UC10-4B9 | Biolegend |
| | CD45RB | C363-16A | Biolegend |
| | H2 | M1/42 | Biolegend |
| | IFNγ | XMG1.2 | Invitrogen |
| | H-2Kb-SIINFEKL | 25-D1.16 | Biolegend |
| PECF594 | TIM3 | B8.2C12 | Biolegend |
| PE-Dazzle 594 | CD11b | M1/70 | Biolegend |
| | CD69 | H1.2F3 | Biolegend |
| PE-Cy7 | CD49b | DX5 | Biolegend |
| | CD62L | MEL-14 | Biolegend |
| | CD86 | GL-1 | Biolegend |
| | IL-2 | JES6-5H4 | Invitrogen |
| | CD11b | M/70 | Biolegend |

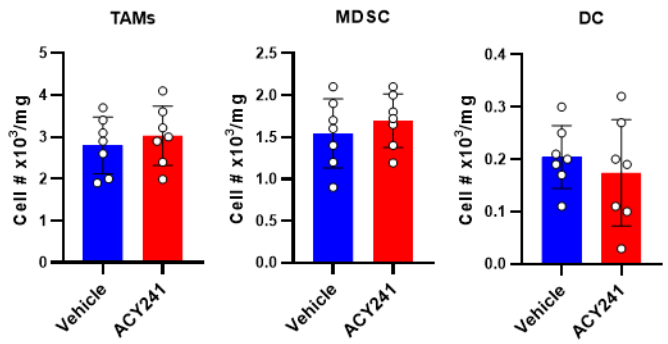


**Fig. S1. Proportion of myeloid cell subsets in the tumors of ACY241- treated lung cancer-bearing mice.** Tumor nodules harvested from KP mice treated for 3 weeks with vehicle or ACY241 were analyzed by flow cytometry to assess the proportions of immune cell subsets. Shown is cell quantity per milligram of tumor for tumor-associated macrophages (TAMs; CD11b+CD11c-Gr1-), Myeloid-derived suppressor cells (MDSC; CD11b+Gr-1+), and dendritic cells (DC; CD11c+CD11b-) in the tumors of mice treated as indicated.

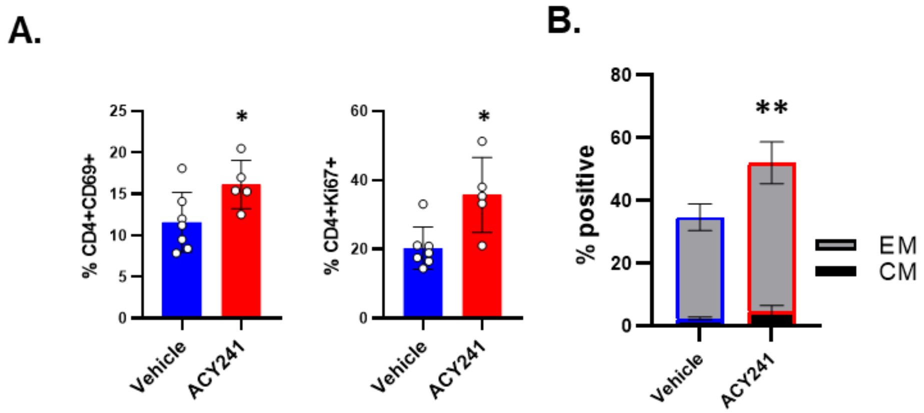


**Fig. S2. Increased activation status of CD4+Foxp3- T cells in the tumors of ACY241- treated lung cancer-bearing mice.** The phenotype of tumor-infiltrating CD4+Foxp3- T cells in the tumors of KP mice treated for 3 weeks with vehicle or ACY241 was assessed by flow cytometry. (A) Expression of CD69 (left) and Ki67 (right). (B) Percent of the CD4+Foxp3- T cells with an effector memory (EM) or central memory (CM) phenotype within tumor-infiltrating CD45+CD3+ cells as determined by CD62L and CD44 staining. Data in (B) are mean ± SEM of 5-7 mice per group. * indicates p-value < 0.05, ** p-value <0.01.

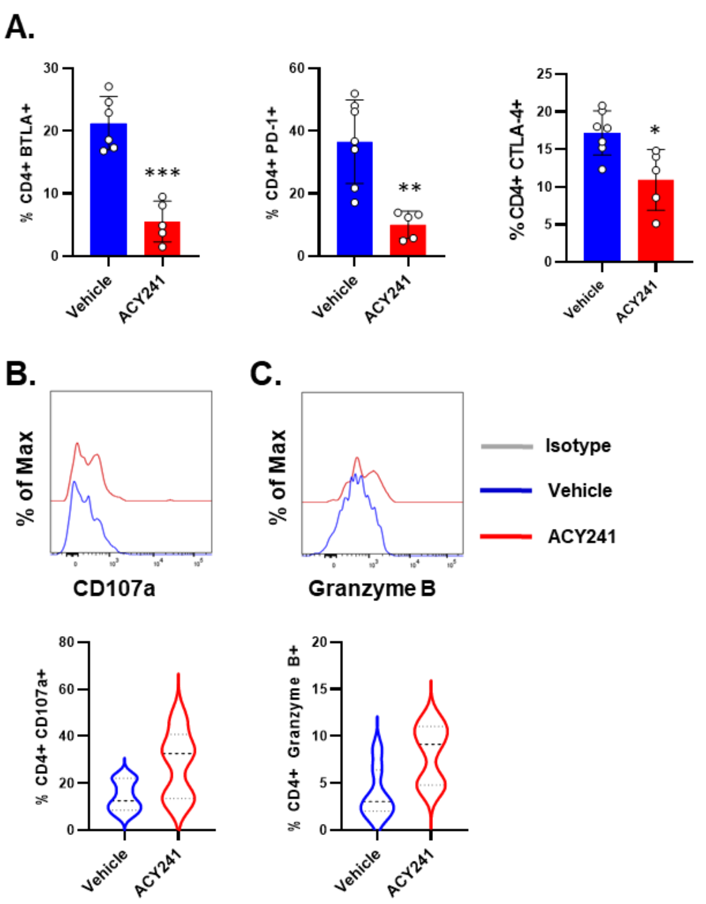


**Fig. S3. Increased effector signature in tumor-infiltrating CD4+Foxp3- T cells of ACY241-treated mice.** Single cell suspensions generated from lung tumors of KP mice that were treated with vehicle or ACY241 were analysed by flow cytometry to determine the phenotype of tumor associated CD45+CD3+ T cells. (A) Expression of BTLA (left), PD-1 (middle) and CTLA-4 (right) on tumor-infiltrating CD4+Foxp3- T cell subset. (B, C) T cells isolated from from lung tumors of treated mice were stimulated for 6 hours with PMA+ Ionomycin in the presence of golgi inhibitor for subsequent intracellular cytokine staining. Representative histograms (top) and summary (bottom) for the expression of CD107a (B) or granzyme B (C) on gated CD4+Foxp3- T cells. Data in B and C (bottom) are mean ± SEM of 5-7 mice per group. * indicates p-value < 0.05, ** p-value <0.01, *** p-value <0.001.

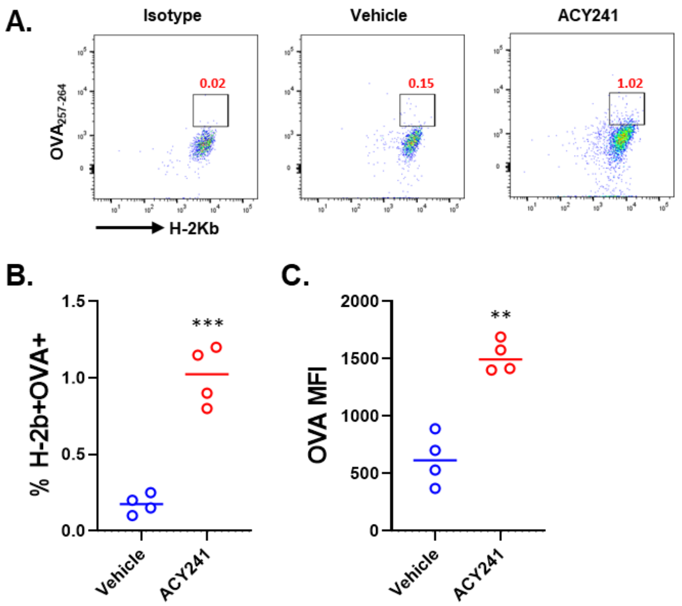


**Fig. S4. Increased expression of MHC class I-associated tumor antigen on TAMs upon ACY241 treatment.** B6 mice were orthotopically implanted with KP tumor cell line that expresses both CD4-specific ($OVA_{323-339}$) and CD8-specific ($OVA_{257-264}$) OVA epitopes (B6 OVA 10103 F LT1 OVA-PGKKP). Mice were treated for 2 weeks with either vehicle or ACY241 after which tumor nodules were processed into single cell suspensions and analyzed for the expression of $OVA_{257-264}$ associated with MHC class I, H-2Kb on CD11b+CD11c-Gr1- TAMs. (A) Representative dot plots for the expression of H-2Kb and $OVA_{257-264}$ in ACY-treated tumors stained with isotype antibody (left), or tumors of vehicle (middle) and ACY241 (right)

treated mice stained with anti $OVA_{257-264}$ antibody. (B) Summary of H-2Kb+ $OVA_{257-264}$+ gate on TAMs. (C) Summary of OVA expression on CD45-EpCAM+ tumor cells based on median fluorescent intensity (MFI). Data in A is representative of 4 mice per group. ** indicates p-value <0.01, *** p-value <0.001.

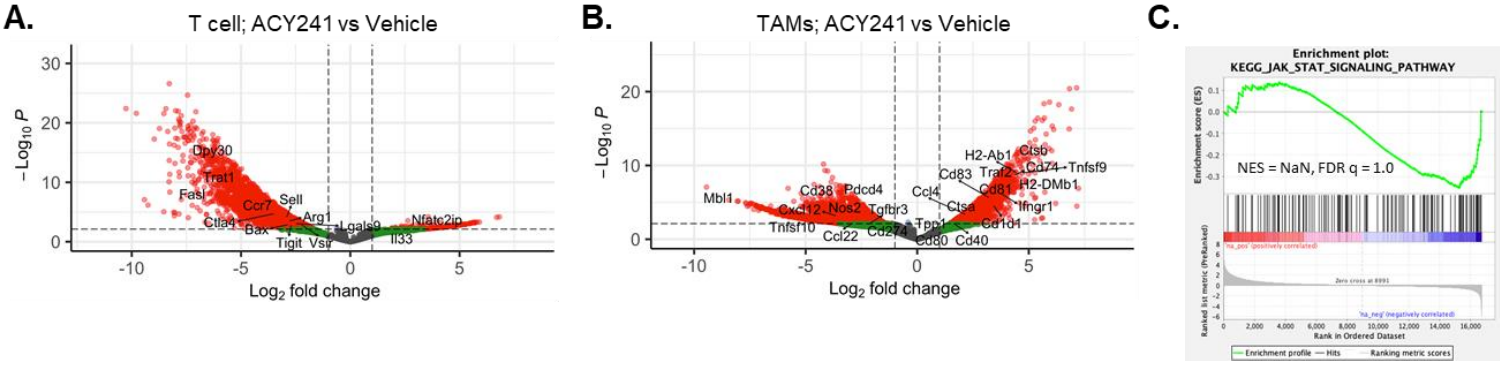


**Fig. S5. Gene expression profile for tumor-associated T cells and macrophages in KP mice treated with ACY241** Transcriptome changes in CD3+Foxp3- T cells and CD45+CD11b+CD11c-Gr-1- (TAMs) that were sorted from lung tumors of vehicle and ACY241-treated KP mice were evaluated by bulk RNA sequencing. Volcano plot for the differential expression of gene transcripts in T cells (A) and in TAMs (B) expressed as Log2 Fold Change in ACY241-treated samples relative to vehicle controls. Padj <0.05 was used as cut-off to determine significance. Gene set enrichment analysis (GSEA) was performed to identify pathways highly enriched in TAMs from ACY241-treated mice over vehicle. (C) GSEA plot for JAK/STAT signaling pathway.

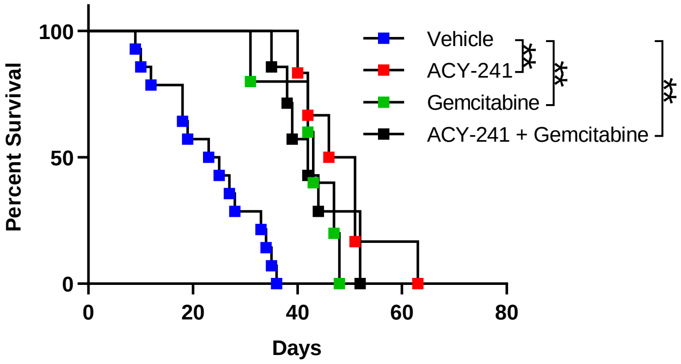


**Fig. S6. Lack of therapeutic benefit under ACY241 and Gemcitabine combination in lung tumor-bearing mice.** KP mice with established tumors (150 – 200mm$^3$) received intraperitoneal injections of ACY241 (3x/week) and/or Gemcitabine (1x/week) for 6 consecutive weeks. Mice receiving vehicle served as controls. Shown is kinetics of survival for tumor-bearing mice that were monitored until clinical endpoint. Data is from 5-7 mice per treatment group. ** indicates p-value <0.01.

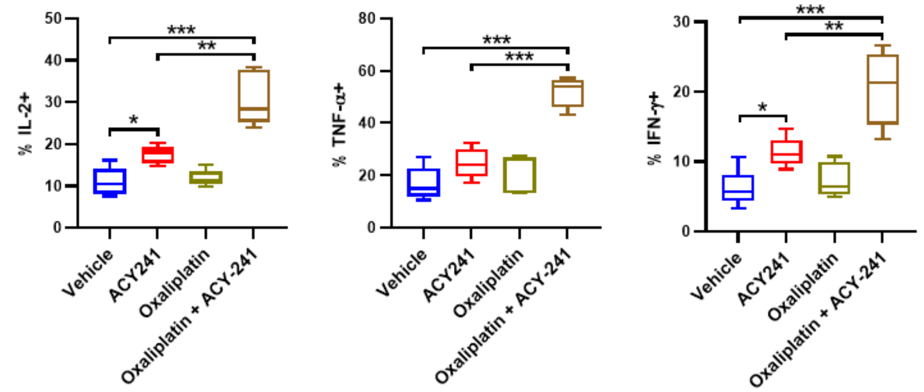


**Fig. S7. Increased capacity for effector cytokine secretion by tumor-infiltrating CD4+ T cells after ACY241 and Oxaliplatin treatment of lung tumor-bearing mice.** KP mice treated with ACY241 and/or Oxaliplatin for 6 weeks were monitored until clinical endpoint at which time tumors were resected. Tumor-infiltrating CD45+CD3+ T cells were then isolated from the tumor cell suspensions and stimulated for 6 hours with PMA+ Ionomycin in the presence of Golgi inhibitor for subsequent intracellular staining. Percent of gated CD4+ T cells that produced IL-2 (left), TNF-α (middle), and IFNγ (right). Data are mean ±SEM of 5-7 mice per group. * indicates p-value < 0.05, ** p-value <0.01. *** p-value <0.001.

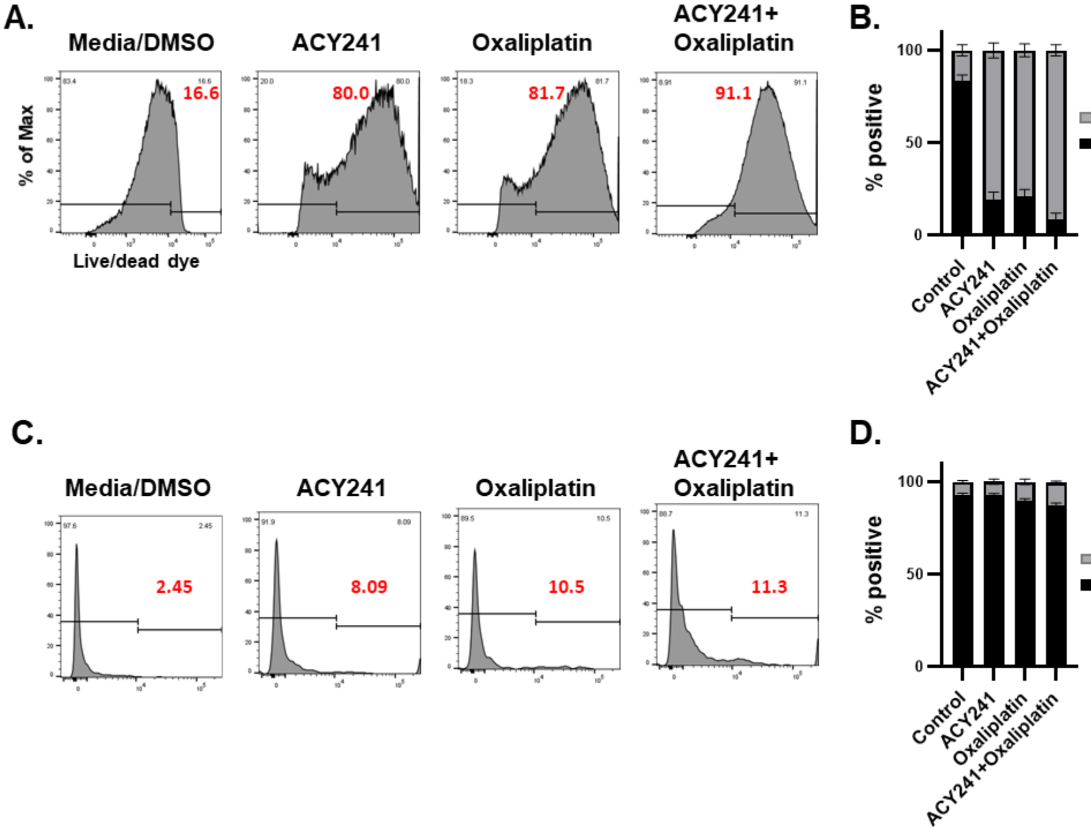


**Fig. S8. Lack of overt toxicity on tumor-associated immune cells treated with ACY241 in combination with Oxaliplatin.** Freshly resected non-small cell lung cancer patient tissues were processed into single cell suspensions and cultured in complete media supplemented with low dose IL-2 (10IU/ml) and IL-15 (10ng/ml) in the presence of ACY241 and/or Oxaliplatin for three days. DMSO in media was added to control cultures at similar concentration (0.01%) as the drug formulations. Staining with live/dead viability dye was performed to determine live and dead cells in EpCAM+ tumor cells or CD45+ immune cells. (A, C) Representative histograms for proportion of tumor (A) or immune (C) cells that are dead as determined by positive staining for the live/dead viability dye. (B, D) Summary for the proportion of live versus dead cells in EpCAM+ tumor (B) or CD45+ immune cells (D). Data are representative (A, C) or mean ±SEM (B, D) of 3 independent experiments.

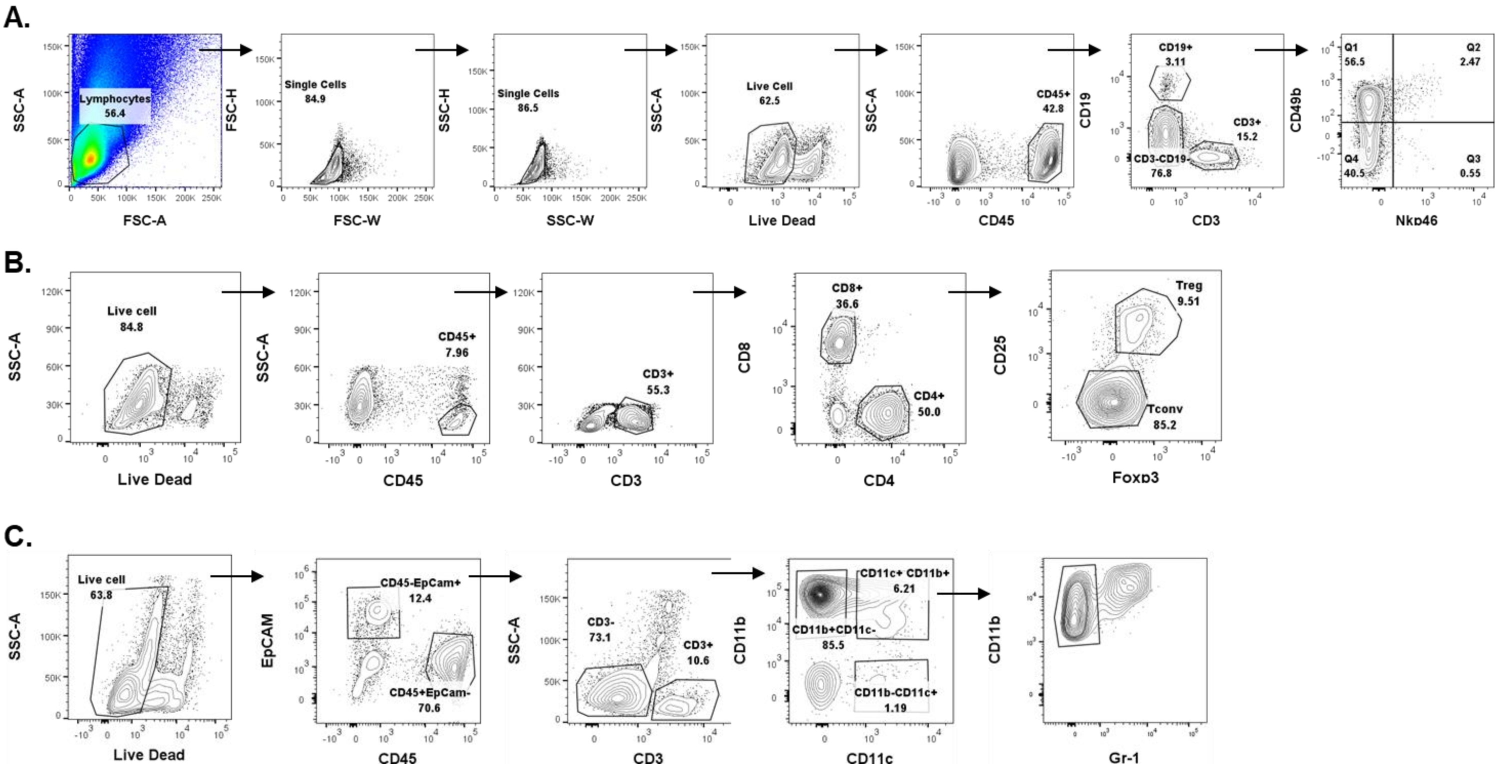


**Fig. S9. Gating strategy for flow cytometry analysis.**

Gating strategy for (A) NK cells, (B) T cell subsets, and (C)Myeloid cell subsets.
