## Supplementary material for "The immunomodulatory properties of the HDAC6 inhibitor ACY241 supports robust anti-tumor response in NSCLC when coupled with the chemotherapy drug Oxaliplatin": Cover letter to JITC

October 1st, 2021

Dear Editor,

Enclosed is our manuscript entitled "The immunomodulatory properties of the HDAC6 inhibitor ACY241 supports robust anti-tumor response in NSCLC when coupled with the chemotherapy drug Oxaliplatin" which we hope will be considered for publication in the Journal for Immunotherapy of Cancer.

This study focuses on understanding the scope of the immunomodulatory effect of ACY241 in NSCLC. ACY241 a second-generation, orally available HDAC6 inhibitor that is structurally similar to ricolinostat, but which exhibits better solubility profile. In this report, we found that it evokes a profound effect on tumor-associated T cells, NK cells, and macrophages in a manner that is more expansive than what has been previously reported for ricolinostat (Adeegbe, et. al., Cancer Discov. 2017 August; 7(8): 852–867). Striking transcriptional changes in tumor-associated T cells and macrophages also accompanied administration of this HDAC6 inhibitor providing further insight into how it likely potentiates its immunomodulatory effects in the setting of NSCLC.

Impressively, ACY241 promoted anti-tumor response as a single agent in our pre-clinical mouse model of Non-small cell lung cancer (NSCLC), an effect that was robustly amplified when coupled with the chemotherapy agent Oxaliplatin. While ACY241 has been tested in early clinical trials with other agents like anti-PD-1 or paclitaxel, evidence for its robustness to support therapeutic effect in NSCLC remains sparse. It's in this regard that our study will be of broad appeal to the JITC readership as it provides compelling observations for the immunogenomic effects of ACY241 and provides rationale for clinical testing of ACY241 and Oxaliplatin as a novel immunotherapy drug regimen for NSCLC therapy. Given the immunomodulatory effects ascribed to ACY241 as described in the manuscript, we opine that it may spur further interest in exploring its partnership with other agents in other solid cancers or even in pathological conditions in which leveraging the described effects could potentiate therapeutic outcomes.

We would like to suggest the following potential reviewers for our study given their relevant expertise:

1. Dr. David Woods
Assistant Professor
Department of Medical Oncology
University of Colorado Denver – Anschutz Medical Campus


2. Jeffrey S. Weber
   Deputy Director, Perlmutter Cancer Center
   NYU Langone Health
  

3. Mark Awad
   Assistant Professor
   Department of Medical Oncology
   Clinical Director, Lowe Center for Thoracic Oncology
   Dana-Farber Cancer Institute
   Harvard Medical School
  

4. Joseph Barbi
   Assistant Member
   Department of Immunology
   Roswell Park Comprehensive Cancer Center
  

5. Giuseppe Giaccone
   Professor and Director, Thoracic Medical Oncology
   Department of Medicine
   Meyer Cancer Center at Weill-Cornell Medical University
  

6. Dmitry Gabrilovich
   Chief Scientist, Cancer Immunology
   AstraZeneca
  

We hope that you will find our study suitable for peer-review consideration in JITC and look forward to your guidance in this process.

Sincerely,

**Dennis O. Adeegbe, Ph.D.**
Assistant Member, Department of Immunology
Assistant Member, Department of Thoracic Oncology
H. Lee Moffitt Cancer Center
Assistant Professor, Department of Oncologic Sciences, USF
